# Decision uncertainty as a context for motor memory

**DOI:** 10.1101/2023.03.15.532761

**Authors:** Kisho Ogasa, Atsushi Yokoi, Gouki Okazawa, Morimichi Nishigaki, Masaya Hirashima, Nobuhiro Hagura

## Abstract

The current view of perceptual decision-making suggests that once the decision is made, a single motor program associated with the decision is carried out, irrespective of the degree of uncertainty involved in the decision-making process. As opposed to this view, we show that different levels of decision uncertainty contextualize actions differently, allowing the brain to form different motor memories based on each context. The match between decision uncertainty during learning and retrieval is critical for successful motor memory retrieval. The same movement trajectory can be associated with different motor memories if each memory is linked to a different level of decision uncertainty. Encoding motor memories based on decision contexts may enhance the robustness of control during the varying neural activities induced by different cognitive states.

## Introduction

In a penalty shoot-out of a football (soccer) game, one may decide to kick the ball to the right corner confidently, seeing that the goalkeeper is moving to the other side, or decide to make the same kick while being unsure about the goalkeeper’s movement. Because both actions are *apparently* identical, we tend to believe that the same motor memory (i.e., a motor program for kicking the ball to the right) is retrieved and executed for both cases regardless of the quality of the preceding decision. But is this true?

Previous perceptual decision-making studies have treated uncertainty as a factor for modulating the evidence accumulation process for decisions (*1, 2*) (*3*), implicitly assuming that an identical motor program is triggered once the evidence level reaches a bound. However, learning or performing an action differently based on decision uncertainty seems sensible because subjective uncertainty can be correlated with important behavioral factors, such as the expected outcome of an action or the possibility of revising a motor plan (*4*) (*5*).

Here, we show that actions that follow certain and uncertain decisions are encoded and memorized differently. In other words, we demonstrated that decision uncertainty works as a contextual cue for motor memory. This finding contrasts sharply with the dominant view in the field, which postulates that contextual cues for motor memories consist of factors that are directly relevant for motor execution, such as the visual appearance of an object to act on that implies different control dynamics, type or location of reach targets (*6, 7*), and posture/state of other body parts during a certain action (*8, 9*) (*10*). We demonstrate that covert internal decision processes, without involving any other bodily movements, could also be a contextual cue for motor memory.

## Results

### Retrieval of motor memory is tuned to the trained decision uncertainty level

First, we tested whether the action learned under a particular decision uncertainty can be retrieved better when the same decision uncertainty level precedes it. Previous studies on episodic memory have established that the shared context between learning and retrieval facilitates the successful recall of memory (*11*). Therefore, we can predict a similar phenomenon if decision uncertainty can function similarly as a contextual cue for motor memory.

Participants (N=38) judged the direction (left or right) of a visual random-dot motion stimulus presented on a screen (Fig. 1**A**, fig.S1). Participants were assigned to one of two groups during the learning phase. The certain-decision group (n=19) judged the direction of a 100% coherent random-dot motion, whereas the uncertain-decision group (n=19) judged the direction of a 3.2% coherent motion. Following this decision, they made a straight center-out reaching movement towards the target in the direction of the perceived motion (Fig. 1**A**). In the learning phase, a velocity-dependent curl force field (*12, 13*) was applied to the movement. The participants had to make a straight movement by resisting the perturbing force (Fig. 1**B, C**). The force-field trials were interleaved with probe trials, where the action was performed following the random-dot motion decision with different uncertainty levels (probe trials: ±3.2%, 6.4%, 12.8%, 25.6%, 51.2%, and 100% motion coherence levels). The trajectory of reaching during the probe trials was constrained to a straight path between the home position and target (channel), and the force the participants applied to the wall of the channel was measured (error-clamp trials) (Fig. 1**B**). This allowed us to measure the amount of force retrieved and applied to resist the perturbation while avoiding the occurrence of any kinematic errors (*14*); thus, the retrieval of motor memory evoked by different decision contexts can be inferred. If the decision uncertainty preceding the action works as a contextual cue for the motor memory, we can predict the best retrieval performance at the level of decision uncertainty in which the motor memory is formed.

**Fig. 1.**
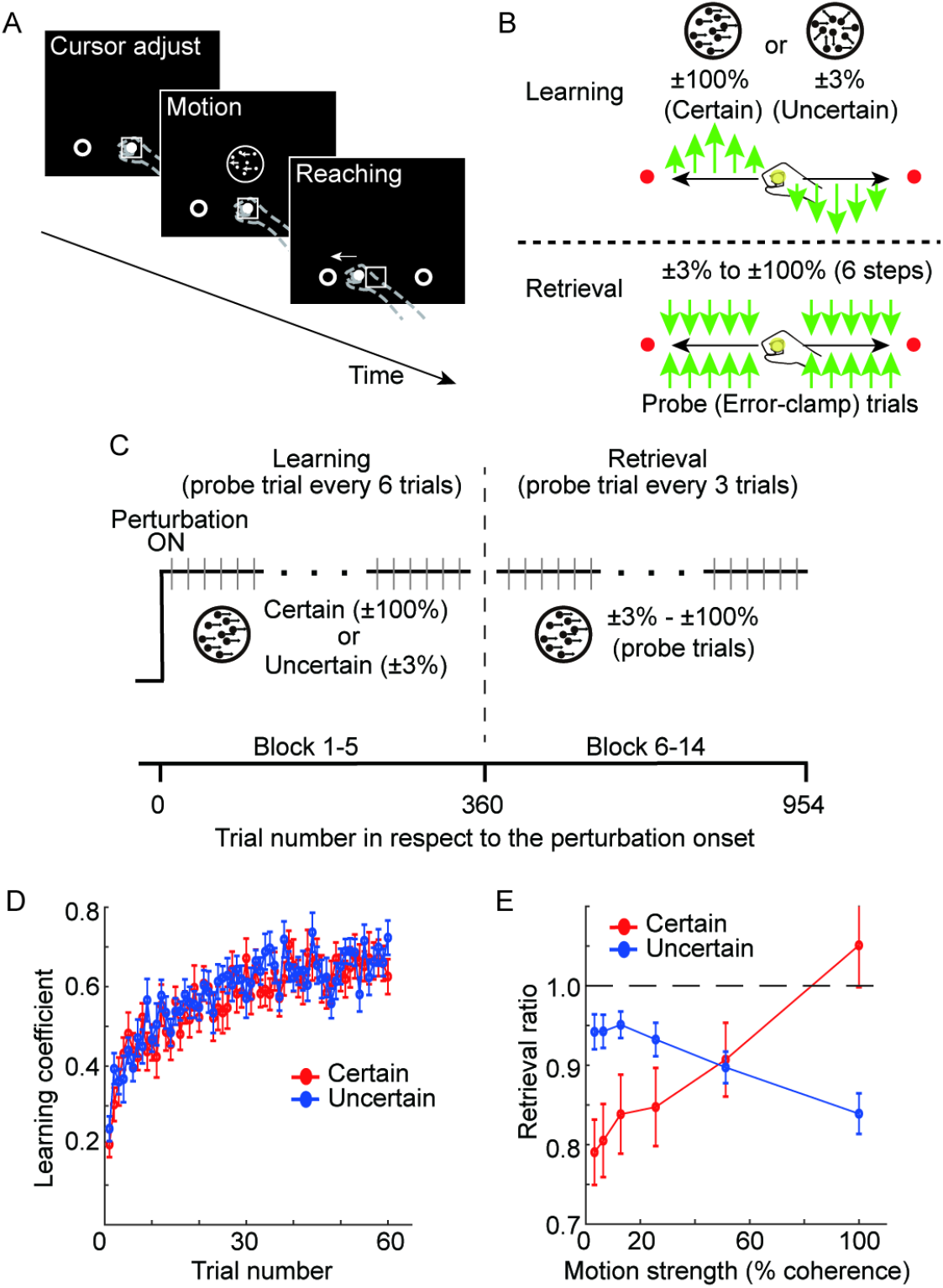
Decision uncertainty-level dependent tuning of motor memory retrieval. **A**: Target-reaching task preceded by motion discrimination. Participants held a handle and judged the direction of a random-dot motion. Their decision was indicated by moving a robotic manipulandum towards the target in the direction of the perceived motion. **B, C**: Force-field learning under different uncertainties. In the learning phase (upper panel in **B**, left in **C**), participants learned to make a straight-reaching movement to the target by resisting a velocity-dependent force field (green arrows). One group learned the force following the decision of a certain stimulus (Certain group;100% coherent motion), whereas another group learned the force following an uncertain stimulus (Uncertain group; 3% coherent motion). In the retrieval phase (lower panel in **B**, right in **C**), both groups of participants performed the task while judging the motion with six different uncertainty levels. The level of retrieval was measured as the amount of force produced against the wall during the error-clamp probe trials (grey lines in **C**), in which the movement trajectory of the hand was constrained to a straight path (error-clamp) (lower panel of **B**). **D**: Progression of force-field learning (probe error-clamp trials) during the learning phase. The vertical axis indicates the leaning coefficient; the value of 1 indicates the full compensation of the perturbation. **E**: Generalization of motor memory across different uncertainty levels in the retrieval phase. The vertical axis indicates the amount of force divided by the amount of force learned at the end of the learning phase (a value of 1 indicates the full retrieval of the motor memory).

Following our prediction, we found distinct retrieval patterns of motor memory between the two groups (Fig. 1**E**). For the certain-decision group, when the motion coherence level was 100% in the retrieval phase, participants were able to produce the learned level of force. However, for the trials with a 3.2% coherent motion, the force dropped to 80% of the learned level (paired t-test, t[18]=8.43, *p*=1.16×10^−7^, dz=1.93). Similarly, for the uncertain-decision group, the same level of force in the learning phase was maintained following a 3.2% motion stimulus. Still, the force level again dropped following the decision of 100% coherent motion (paired t-test, t[18]=5.53, *p*=2.98×10^−5^, dz=1.27). Thus, the manner in which the force was retrieved depended on the decision uncertainty level at which participants learned the force field (Fig. 1**E**; analysis of variance [ANOVA] interaction effect; F[5,180] = 61.46, *p*=3.91×10^−37^, η^2^=0.63). Such reversed retrieval patterns of force between the two groups cannot be explained by the generally deteriorated motor output following uncertain decisions (*15*) since this would predict that the force would drop towards higher-uncertainty decisions regardless of the different learning experiences.

Furthermore, the difference in motor-learning quality or decision-making performance between the two groups cannot explain this result. First, the rate and magnitude of motor learning were comparable between the two groups (Fig. 1**D**). Second, both groups showed higher accuracy and faster reaction times for higher motion coherence during the retrieval phase, as expected (fig. S2). The overall correct rates and reaction times were moderately higher for the uncertain-decision group, probably due to the differences in the speed-accuracy trade-off (*16*) and effect of perceptual learning with different task difficulties (*17*). Still, such differences do not explain why the two groups produced opposite force production patterns. Rather, the result suggests the independence of action initiation and the quality of action execution (*18*), which the latter reflects the retrieved content of motor memory.

Finally, it is unlikely that the feature of the visual stimulus (100% and 3% coherent motion) is the main determinant of this effect since visual cues on their own are known to be weak contextual cues for the retrieval of motor memory (*19*) (see also fig. S3 for the control experiments).

Taken together, the result of the incomplete transfer of motor memory across different decision uncertainties implies that the decision process preceding the action can be a context for the motor memory.

### Motor memory can be tagged by different decision uncertainty contexts

A more direct test for context-dependent motor learning is to show that participants can simultaneously learn two different force fields associated with different contexts for the same reaching movement (*8, 19*). In Experiment 2, using a within-participant design, we directly examined whether decision uncertainty can indeed function as a contextual cue for tagging different motor memories, enabling the learning of two different force fields.

In this study, 100% and 3.2% were used as the coherence levels of the motion stimulus. After the baseline phase, the participants were exposed to two different force fields (Fig. 2**B**): The strong and weak force fields for Experiment 2-1 (n=19; Fig.2**A** upper panel) and two opposing force fields (clockwise [CW] or counterclockwise [CCW]) for Experiment 2-2 (n=18, Fig.2**A** lower panel). In both experiments, two different decision uncertainties were associated with either of the two different force fields. Suppose the brain can use decision uncertainty to segregate the context and retrieve the relevant motor memory. In that case, participants should be able to learn and retrieve strong and weak forces (Experiment 2-1) or force in the CW and CCW directions (Experiment 2-2), depending on the preceding decision uncertainty type (certain or uncertain). In contrast, if the difference in the decision process is insufficient to tag different force fields, the output force level after learning would not differ between the two force-field conditions.

**Fig. 2.**
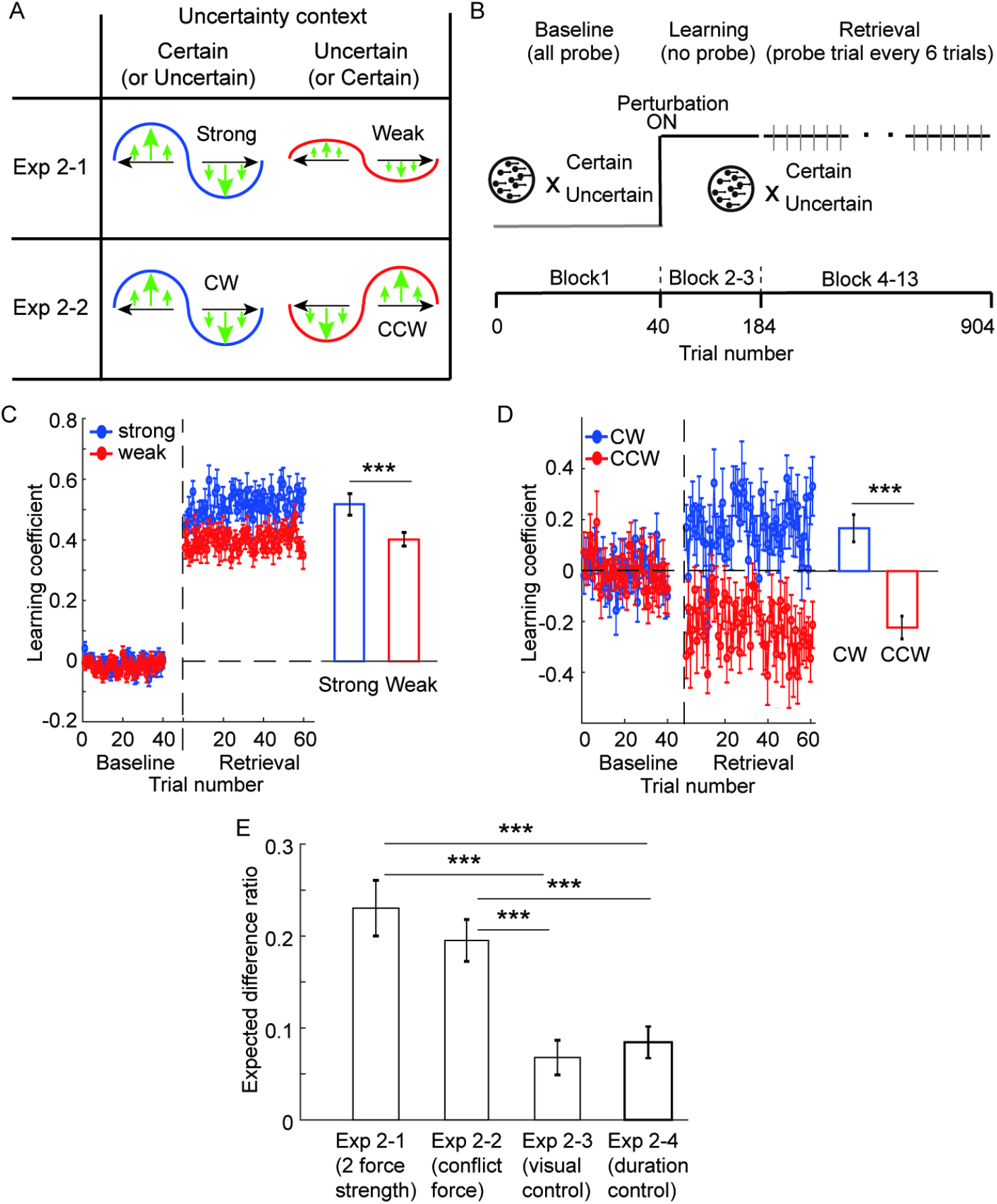
Simultaneous learning of different force fields based on each decision uncertainty context. **A, B**: Conditions of Experiment 2-1 and 2-2 (**A**) and the trial structure of the experiments (**B**). Each uncertainty context, the decision of 100% or 3% coherent motion, was associated with a strong or weak force field in Experiment 2-1 and with opposing force-field directions in Experiment 2-2. The grey line indicates the probe error-clamp trials. **C, D**: Force output measured in probe error-clamp trials during the baseline and the retrieval phase. For Experiment 2-1 (**C**), the data are aligned depending on the strength of the force (strong and weak). Note that the vertical axis shows the learning coefficient for the strong force. Thus, the full compensation of the weak force will be 0.5. For Experiment 2-2 (**D**), data are aligned to the direction of the force (CW and CCW). **E**: Comparison of the effect with the (visual and duration) control experiments, expressed as the ratio to the expected difference (index of contextual learning; see Supplementary Methods). Error bars indicate the standard error of means across participants. ***: *p*<0.001.

This result supports our hypotheses. After the learning, participants produced relevant levels of force associated with the given decision context, producing stronger force for the strong-force condition than the weak-force condition in Experiment 2-1(paired t-test, t[18]=7.63, *p*=4.78×10^−7^, dz=1.75) (Fig. 2**C**) and producing the force in the opposing directions in Experiment 2-2 (paired t-test, t[17]=8.15, *p*=2.85×10^−7^, dz=1.91) (Experiment 2-3; Fig. 2**D**).

With another set of participants, we confirmed that such a difference could not be observed when the random-dot motion with different coherence levels was associated with different force fields but without involving any decision about the motion direction (Experiment 2-3; Fig. 2**E**, fig.S3; see details in the Supplementary Methods). This shows that the results of Experiment 2-1 and 2-2 cannot be simply explained by the difference in the associated visual input pattern itself, corroborating previous literature findings (*19*) (independent t-test; Exp2-1 vs. 2-3; t[37]=4.61, *p*=1.83×10^−4^ [corrected], dz=1.19, Exp2-2 vs. 2-3; t[36]=4.22, *p*=6.28×10^−4^ [corrected], dz=1.13) (see also Experiment 3 results). We also confirmed that the difference in stimulus duration between easy and difficult stimuli could not explain the results (Experiment 2-4; Fig. 2**B** and fig.S3) (Exp2-1 vs. 2-4: t[34]=4.07, *p*=0.0011 [corrected], dz=1.13, Exp2-2 vs. 2-4: t[33]=3.71, *p*=0.0030 [corrected], dz=1.07). Taken together, the results show that preceding decision uncertainty indeed works as a contextual cue for the learning and retrieval of distinct motor memories.

### Decision uncertainty, not the perceptual uncertainty, contextualized motor memory

Finally, we investigated what constitutes this type of novel uncertainty context, whether it is tied to the uncertainty of a specific input stimulus (e.g., random-dot motion) or whether it is a stimulus invariant, abstract uncertainty about the decision. In the latter case, participants should be able to retrieve motor memory even when visual stimuli are different between the learning and retrieval phases, if the uncertainty level is matched.

To examine this, we used two types of visual stimuli in Experiment 3. One is random-dot motion, as was used in the previous experiments (motion stimulus), while the other was an arrow stimulus in which a sequence consisting of left and right arrows was presented in a short period of time (20 arrows in 1,500 ms) (Fig. 3**A**). In the arrow stimulus, participants were asked to decide which of the two stimuli (left or right arrow) was presented more frequently after the termination of the sequence and then immediately reach towards the target in the direction of their decision. Uncertainty was manipulated by changing the ratio of the left and right arrows in the sequence. Before the main experiment, we matched the confidence level of decisions (i.e., subjective estimate of decision uncertainty) between the random-dot motion and arrow stimuli based on the participants’ confidence reports from a separate experiment. Decision confidence of the arrow stimulus with a left-right ratio of 5.5: 4.5 (5% bias from chance) corresponded to a 3% coherence level random-dot motion stimulus. Likewise, a ratio of 9:1 (40% bias from chance) in the arrow stimulus corresponded to the 100% coherence level random-dot motion (Fig. 3**C**).

**Fig. 3.**
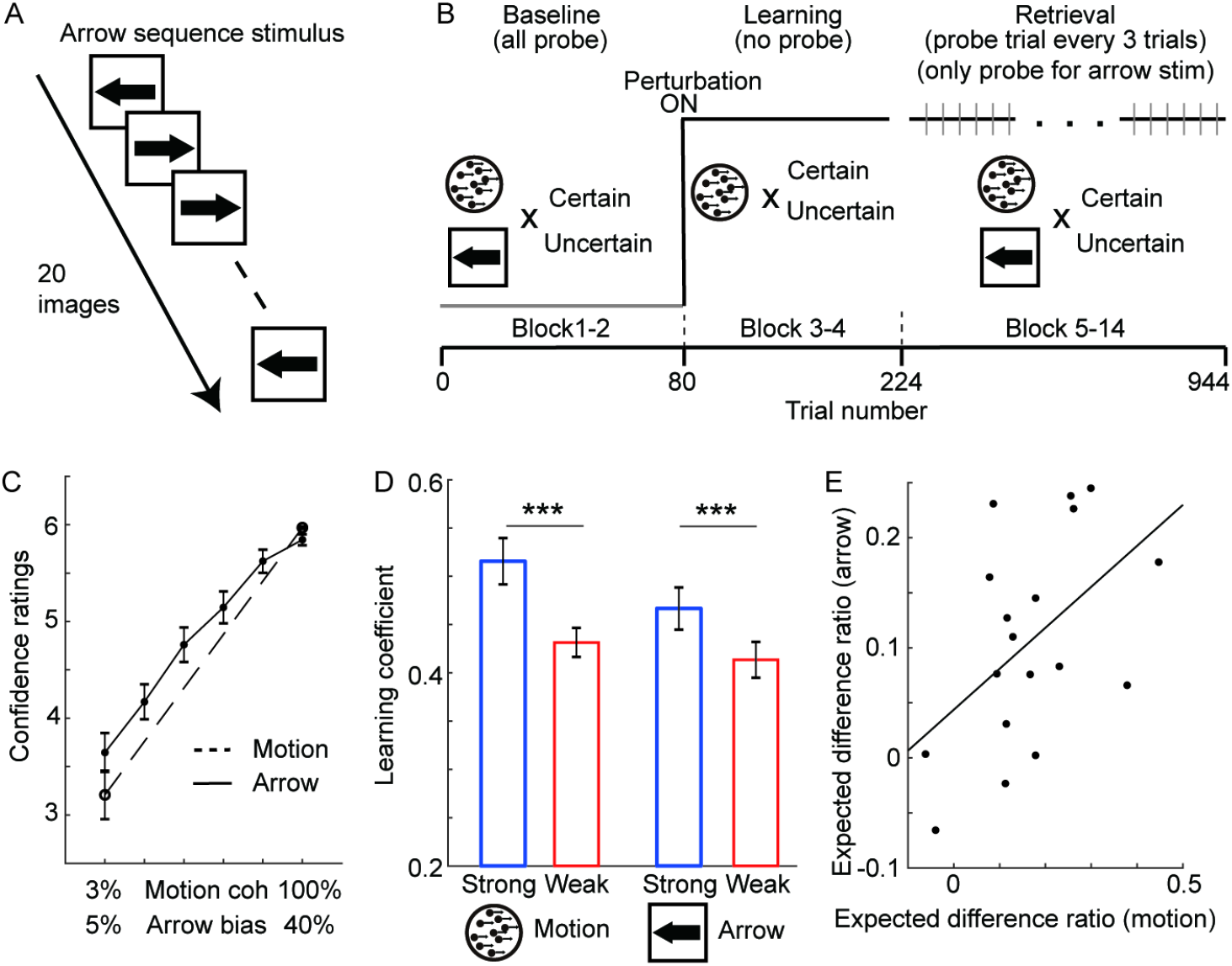
Transfer of decision uncertainty context across different visual stimuli. **A**: Arrow sequence stimulus in Experiment 3. A sequence of 20 arrows was presented, and the participants judged which arrow direction (left or right) had a higher probability in the sequence. **B**: Conditions and the trial structure of Experiment 3. In the baseline phase, participants made motion discrimination or arrow direction decision and reached towards the target without any perturbation. For both visual stimuli, certain and uncertain stimuli were prepared, in which the confidence level was matched across different visual stimuli. In the learning phase, participants learned the force only during the motion direction decision, in which strong and weak force-field was associated with different uncertainty levels (certain and uncertain). In the error-clamp trials of the retrieval phase, together with the certain and uncertain motions, the arrow sequence stimulus was again presented as in the baseline phase. Note that the reaching following the arrow decision in the retrieval phase was only performed for the probe error-clamp trials (grey line). **C**: The match of the confidence level across different types of visual stimulus (random-dot and arrow-sequence). **D, E**: Force retrieved in each condition. The force field tagged with different motion uncertainty levels (left) was able to be retrieved using different visual stimulus (arrow stimulus; right) with similar uncertainty levels (**D**). Across participants, the amount of difference of the force (i.e., index of contextual learning) in the arrow stimulus correlated with that of the motion stimulus (R=0.51, *p*<0.05) (**E**). Note that participants have never experienced the force field following the arrow stimulus decision. Error bars indicate the standard error of means across participants. ***: p<0.001.

As in Experiment 2-1, participants learned two different strengths of force fields (strong and weak) associated with motion stimuli with 100% or 3% coherent motion. After learning, we tested whether arrow stimuli could retrieve the force learned under random-dot motion stimuli with matched confidence levels. Importantly, during the task, all arrow-stimulus trials were error-clamped (Fig. 3**B**). This means that participants never experienced force perturbation while performing the action following the arrow stimulus decision. Therefore, any force produced to resist perturbation during the arrow stimulus is the component transferred from the motor memory formed under the random-dot-motion stimulus.

First, we replicated the results of Experiment 2-1. The participants were again able to learn two different force fields associated with two different uncertainty levels of the motion decision. The amount of force produced during the error-clamp trials was significantly different between the two different force-field conditions (paired t-test, t[17]=5.50, *p*=3.91×10^−5^, dz=1.29) (Fig. 3**D** left, Supplementary Fig. 4). Second, and more critically, for the trials with arrow decisions, we also found a significant difference in the force between the strong and weak force field conditions (paired t-test, t[17]=4.73, *p*=1.92×10^−4^, dz=1.16) (Fig. 3**D** right, fig. S4). Finally, the individual differences in force between the two force fields (i.e., index of contextual learning) were correlated between the random-dot motion condition and arrow-sequence condition (Fig. 3**E**; R[18]=0.51, *p*=0.032), suggesting a shared component between the two variables. These results clearly show that motor memory encoded with random-dot motion can be retrieved using different visual stimuli with similar decision uncertainty levels. In other words, part of the motor memory is tied to abstract decision uncertainty, which is invariant from the feature of the input stimulus.

## Discussion

The context for encoding memory has been of great interest in the field of cognitive neuroscience (*11*) (*20, 21*) (*22*). For the domain of motor memory, the majority of the contexts identified are directly involved in the overt or ongoing motor control process, for example, the spatial position of the workspace (*19*), direction of the planned movement in the workspace (*6*), plan of the future state (*7*) or concurrent state of the relevant or irrelevant body parts (*8, 9*) (*10*). Our study demonstrated that covert internal decision processes, without any overt difference in the bodily state, could also be a contextual cue for motor memory, adding a novel dimension for the contexts to be considered.

Uncertainty about how an action will be perturbed has been shown to impact motor learning, where the learning rate is modulated depending on the stability of the environment (*23*). This phenomenon cannot simply explain our results because the amount of learning itself did not depend on the uncertainty level of the decision (Fig. 1**C**). This indicates that coping with the uncertainty of decisions and coping with the uncertainty of perturbations are governed by different processes in the brain; for the former, the brain contextualizes motor memory depending on decision uncertainty.

During perceptual decision-making, the ongoing state of evidence accumulation during the deliberation period is reflected in neuronal activity in the cortical areas involved in motor planning and execution (*24*) (*25*) (*26*) (*27*) (*28*). Perceptual evidence guides an agent’s decision, but at the same time, the agent can calculate the subjective uncertainty level of the decision (i.e., decision confidence) using accumulated evidence signals (*29*). Indeed, neuronal activities in both cortical (*29*) and subcortical structures reflects the uncertainty of action to perform (*30*) (*31*). We speculate that such premovement activity reflecting decision uncertainty (confidence) forms the *context*, or the neural state, when forming motor memory in the sensorimotor network. Consequently, the action learned in such a context will be best performed (i.e., retrieved) when the same premovement activity pattern is elicited before the action (*32*).

In conclusion, we showed that the brain uses decision uncertainty as a contextual cue to retrieve motor memory, thus preparing different motor memories depending on the uncertainty level of decisions. This indicates that football players should practice not only kicking the ball precisely to the place they want, but also practicing it in both situations when they are sure and unsure about the goalkeeper’s movement.

## Materials and Methods

### Participants

A total of 147 right-handed participants volunteered in Experiment 1 (certain group; 22 [7 women, ages 19–25 years], uncertain group; 22 [7 women, age 20–25 years]); Experiment 2-1 (21 [5 women, age 20–28 years]); Experiment 2-2 (20 [8 women, age 20–30 years]); Experiment 2-3 (20 [7 women, age 20–38 years]); Experiment 2-4 (17 [5 women, age 21–29 years]); and experiment 3 (20 [7 women, age 21–46 years]). All participants were naive to the purpose of the experiment. All experiments were undertaken with the understanding and written consent of each participant following the Code of Ethics of the World Medical Association (Declaration of Helsinki) and with the approval of the National Institute of Information and Communications Technology (NICT) ethical committee. No adverse events occurred during either of the experiments. Experiment 1 used a relatively larger sample size compared to typical motor learning studies because of the cross-subject design. To ensure a similar level of effect size as in Experiment 1, we used a similar number of participants in the rest of the experiments.

### Data and participant exclusion criteria

In each experiment, trials were excluded if the 1) reaction times (movement onset concerning the visual stimulus onset) were too fast (<100 ms; likely not judging the stimulus) or too slow (1,500 ms>; judging after the stimulus disappearance), 2) did not reach properly to the target (<75% of the maximum distance), and when the movement direction reversed after going 2.5 cm to the opposite direction before reaching to the target. If the trial exclusion rate exceeded 30% of the data in the last block of the learning phase or the retrieval/test phase, the participants were excluded from further analysis. Furthermore, if the overall choice rate during the retrieval phase was biased towards one direction (>70%) (e.g., moving [making a decision] to the right in most of the trials), the participant was also excluded because of the asymmetrical motor learning experience between the two directions. See the method section below for task details. Note that these exclusion criteria were set to exclude data/participants who did not follow the instructions of the experiments and maintain the same data quality across participants. However, including excluded participants in the analysis did not qualitatively change the results.

Based on the above criteria, in Experiment 1, three participants from each certain and uncertain group were excluded. Likewise, two participants were excluded from the analysis of Experiment 2-3, 2-4, and 3, respectively.

### General settings

The participants were seated comfortably in front of a screen placed horizontally in front of them, which prevented direct vision of their hands (fig. S1). The visual stimulus was presented on a screen using a projector placed above the screen. The viewing distance was set to 50 cm. The upper trunk was constrained using a harness attached to the chair to maintain the viewing distance. During the experiment, participants were asked to hold the handle of the manipulandum with their right hand (PHANToM Premium 1.5 HF, SensAble Technologies, Woburn, MA, USA), whose position was sampled at 500 Hz. The handle position was displayed as a white cursor (circle, 6 mm in diameter) on a black background on a horizontal screen located above the hand. The movement of the handle was constrained to a virtual horizontal plane (10 cm below the screen) that was implemented by a simulated spring (1.0 kN/m) and dumper (0.1 N/ms^−1^).

The random-dot motion stimulus was presented at the center of the screen (*33*) (*34*) (Fig. 1A). In a 7° diameter circular aperture, dots were presented at a density of 3.5 dot/deg^2^. The speed of the dots is 10°/s. For each trial, either 3.2%, 6.4%, 12.8%, 25.6%, 51.2%, or 100% of the dots moved coherently to the left or to the right (hereafter referred to as motion coherence level). All other dots moved in random directions and were picked for each dot separately between 0° and 360°. The visual stimulus and robotic manipulandum were controlled using an in-house software program developed using C++ (*6*).

Before each trial, the robotic manipulandum automatically guided the participant’s hands to the starting position. A trial started when the participants maintained the cursor at the starting position for 500 ms. Subsequently, a random-dot motion was displayed. Immediately after the decision, participants made a reaching movement either towards the left or right target, depending on their decision (Experiment 1 and 2). The motion stimulus disappeared when the movement was initiated. In Experiment 3, the participants were required to move after the disappearance of the motion stimulus. Each target was located 10 cm horizontal from the starting position.

A velocity-dependent curl force field (*12*) was used for motor learning. The force field was applied according to the following equation:

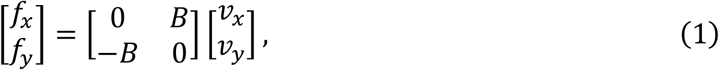

where fx and fy are the forces applied to the handle (N) and vx and vy are the velocities of the handle (m/s) in the x- and y-directions, respectively. For the clockwise (CW) force field, the viscosity coefficient B (N/[ms^−1^]) had positive values, and for the counterclockwise (CCW) field, B had negative values. Channel trials (error-clamp trials) were occasionally introduced to quantify learning of the force field. Here, the handle movement was constrained along a straight path between the home position and target by a simulated damper and spring (*14*), and the force applied to the wall of the channel during the movement was measured. This allowed us to measure the amount of force retrieved to resist the perturbation while avoiding any kinematic errors.

### Experiment 1

We tested how the action learned under a particular decision uncertainty transferred to actions during other levels of decision uncertainty.

#### Procedure

The participants held the handle with their right hands and judged the direction of a random-dot motion (left or right). As soon as they made the decision, they moved their hands towards the target direction corresponding to the direction of the judgment. The random-dot motion disappeared as soon as the participant’s movement was detected (3.5 cm/s). The stimulus disappeared after 1,500 ms, even if no movement was detected (*34*), and participants were instructed to initiate their movement before the disappearance. Before the task, the participants were familiarized with the manipulandum and judgment of the visual stimulus.

The experiment consisted of two phases, learning and retrieval. In both phases, the task was performed under a force field with occasional error-clamp trials (Fig. 1**C**). Half of the participants experienced the CW force field, and the other half experienced the CCW force field. The viscosity coefficient (*B* in Eq. 1) was set to 10 (N/[ms^−1^]).

Participants were divided into two groups: certain and uncertain. During the learning phase, in the certain group, participants learned the force field in response to a 100% coherent motion (low decision uncertainty level). In the uncertain group, participants learned the reaching in response to a 3.2% coherent motion (high uncertainty level). The participants were instructed to maintain the movement trajectory straight, similar to reaching without perturbation. Five blocks of 72 trials were conducted. Error-clamp trials were introduced every six trials between the force-field trials. The motion coherence level during the error-clamp trials was set to be the same as that in the nonerror-clamp trials.

In the retrieval phase, each group of participants performed the same task as that in the learning phase. The only difference was that the frequency of the error-clamp trials was, on average, every three trials, and 12 different coherence levels (+-3.2%, 6.4%, 12.8%, 25.6%, 51.2%, and 100%) were used in these error clamp-trials (positive indicates the direction towards the right and negative to the left). This design allowed us to examine how the motor memory formed at a particular decision uncertainty level generalizes to other levels of uncertainty.

Participants underwent nine blocks, with each block containing 66 trials (22 error-clamp trials; 2 [left and right] trials for 100% coherent motion, 2 trials each for the other 10 motion coherence levels).

### Experiment 2-1

To directly demonstrate the role of decision uncertainty as a contextual cue for motor memory, we tested whether participants could learn two different force fields for the same movement trajectory if each force field was associated with different decision uncertainty levels.

#### Procedure

As in Experiment 1, the participants judged the direction of a random-dot motion and moved the handle towards the target in the judged direction. Two different motion coherence levels, 100% (certain decision) and 3.2% (uncertain decision) were prepared. After the practice session, in the baseline phase, participants performed the task that was error-clamped (two blocks of 40 trials). In the learning phase, participants performed the task under two different strengths of force fields (*B* = 10 [strong] and *B* = 5 [weak] [N/(ms-1)]). Each strong and weak force field is associated with a different preceding decision uncertainty (certain or uncertain). The pattern of the association between the force field strengths, decision uncertainties, and direction of the force fields (CW or CCW) was counterbalanced across participants. The participants underwent two blocks of 72 trials each. In the retrieval phase, participants performed 10 blocks of the task (72 trials) with interleaved error-clamp trials (every six trials; six trials each for two visual stimuli per block).

### Experiment 2-2

We tested whether two force fields in opposing directions could be simultaneously learned if each field was associated with different decision uncertainty levels.

#### Procedure

The setting of the experiment was identical to Experiment 2-1. Still, instead of using strong and weak force fields, we associated two force fields with opposing directions (CW and CCW) with different decision uncertainties (100% and 3.2%). In addition, the participants underwent three blocks of 72 trials during the learning phase. The viscosity level was set to ±2.5 (N/[ms^−1^]) for the CW and CCW conditions.

### Experiment 2-3

As a control experiment, we examined the contextual effect of visual features (100% and 3.2% coherent random-dot motion), which covaried with the decision uncertainty in Experiment 2-1 and 2-2.

#### Procedure

The setting of the experiment was like Experiment 2-1, but the participants were not required to make any directional decision of the random-dot motion. Instead, they either saw 100% or 3.2% coherent random-dot motion presented on the screen. Immediately after the disappearance of the motion stimulus, a single target appeared on either the left or right side, and the participants reached towards the target. The target direction did not correlate with the direction of motion. Thus, the direction of the participant’s movement and decision was unrelated, which discouraged the participants from making decisions in any direction. Duration of the visual stimulus was drawn from the normal distribution, which the mean and the variance were extracted from the reaction times (RT: stimulus onset to the movement onset) in Experiment 2-1 (fig. S2B; parameters: 3.2% motion: 461.9±75.8 ms [left], 453.9±69.9.2 ms [right], 100% motion: 731.5 ±144.3 ms [left], 720.3±149.2 ms [right]). To ensure that the participants focused on the stimulus, they were occasionally asked if the visual motion they saw was coherent or random (12 trials per block). The average correct rate was 91.4±12.7%.

All the other trial structures were identical to Experiment 2-1. After the baseline condition (two blocks of 40 trials; all error-clamped), in the learning and retrieval phases, each coherence level of random-dot motion was associated with either the strong or weak force field in each participant (learning phase: two blocks of 72 trials, retrieval phase: 10 blocks of 72 trials [one error-clamp every six trials]).

A comparable level of force-field learning as in Experiment 2-1 should be observed if the visual feature of the stimulus can be a context for encoding/retrieval of motor memory.

### Experiment 2-4

For another control experiment, we examined the contextual effect of time-before-execution, which also covaries with the decision uncertainty level in Experiment 2-1 and 2-2.

#### Procedure

Setting of the experiment was like Experiment 2-1, but they only observed 100% coherent random-dot motion. Two durations were prepared, in which one corresponded to the RTs (stimulus onset to the movement onset) of 100% coherent motion (short duration) and another to the RTs of 3.2% coherent motion (long duration) in Experiment 2-1. As in Experiment 2-3, this duration was drawn from a normal distribution, in which the mean and variance were extracted from the RTs of the corresponding conditions in Experiment 2-1 (see above).

In this experiment, participants judged the direction of the visual stimulus, reported the decision immediately after stimulus termination, and then made the reaching towards the target in the direction of the judgment. The other parameters were similar to Experiment 2-1. After the baseline phase (two blocks of 40 trials; error clamped), retrieval (two blocks of 72 trials), and test (10 blocks of 72 trials; error-clamp, once in six trials) phases, short and long durations were associated with weak and strong force fields.

Unlike Experiment 2-3, where participants were uninformed of the movement direction until the disappearance of the random-dot motion, this experiment allowed participants to prepare the movement for a longer duration when the stimulus duration was longer. If the stimulus duration and amount of motor preparation are the main components of the context in Experiment 2-1, we should observe an effect comparable to Experiment 2-1 in this experiment.

### Experiment 3

We examine the content of the decision-making uncertainty context. Specifically, we tested whether the uncertainty context includes the abstract stimulus-independent component, other than the input stimulus-level uncertainty, for motor memory retrieval.

#### Procedure

Two different types of visual stimuli were prepared: random-dot motion and arrow sequence. For random-dot motion, participants judged the net direction (left or right) of the dot motion. The uncertainty of the decision was controlled by changing the %-coherence of the dot motion direction. The arrow stimulus consisted of a stream of arrows heading either to the left or right (Fig. 3**A**). A total of 20 arrows were presented in a sequence, each presented for 33.3 ms, followed by a 33.3 ms of the blank. The participants judged the direction of the arrow, which was more frequently presented in the sequence. The uncertainty of the decision was manipulated by changing the left-right ratio of the arrows in the sequence.

#### Matching of subjective uncertainty level (confidence) across the stimuli

First, we established a correspondence in the subjective uncertainty level (i.e., confidence) between the two stimuli. In a trial, either the random-dot motion stimulus or the arrow stimulus was presented for 1,500 ms and then disappeared. After the disappearance of the stimulus, the participants moved the manipulandum towards the target in the direction of their judgment, and no perturbation was applied to this movement. After moving their hand to the target, participants reported the confidence level of the decision on a scale of 0–6, with 0 corresponding to a total guess and 6 corresponding to maximum confidence in the decision. Participants performed five blocks of 64 trials. For the random-dot motion stimulus, two motion coherence levels (100% and 3%) were prepared. For the arrow stimulus, the left-right ratios in the arrow sequence were 55%, 60%, 65%, 70%, 80%, and 90% (5–40% bias). Each block contained 16 random-dot motion stimuli and 48 arrow stimuli.

#### Testing transfer of motor memory across different stimuli

In the confidence matching experiment, we found that the decision confidence for the 5% biased arrow sequence corresponded to the confidence of 3% coherent random-dot motion. Similarly, a 40% biased arrow sequence corresponded to the 100% coherent random-dot motion. Using these four confidence-matched stimuli, we tested the transfer of uncertainty-tagged motor memories across different visual stimuli.

In the baseline phase, all four types of stimuli were presented, and participants underwent two blocks of 40 trials (all error-clamped) (Fig. 3**B**). Next, in the learning phase, only the two coherence levels of random-dot motion (100% and 3%) were presented, in which each was associated with either strong or weak force fields, as in Experiment 2-1. The participants performed two blocks of 72 trials. Finally, all four types of stimuli were presented in the retrieval phase. Here, random-dot motion stimuli had both force and error-clamp trials, but for the arrow stimuli, there were only error-clamp trials. This prevented any learning of force for the arrow stimulus trials, allowing us to purely evaluate the component transferred from learning using a random-dot stimulus. Participants underwent 10 blocks of 72 trials (error-clamp trials; once every three trials).

If the uncertainty context includes the abstract, stimulus invariant component, the motor memory tagged by decision uncertainty of random-dot motion should be retrieved when the arrow stimulus with a matched uncertainty level is presented.

### Data analysis

#### Data analysis of Experiment 1

To analyze the error-clamp trials, the amount of force against the wall of the channel at the timing of the peak movement velocity (velocity peak point) was extracted. Then, the force was divided by the velocity peak value to transform the value into viscosity space (N/[ms^−1^]). Finally, this value was divided by the viscosity of the force field to calculate the % ideal of the force, which represents learning (learning coefficient).

The learning coefficient was calculated for each motion coherence level (collapsed left and right motion data). To assess the generalization of motor memory across different uncertainty levels, the learning coefficient for each motion coherence level was divided by that calculated using the last block of the learning phase (retrieval ratio). Here, a value of 1 represents full retrieval of the memory, and 0 represents complete forgetting.

To quantify the differences in the decision-making process between the certain and uncertain groups, we fitted a drift-diffusion model (DDM) to the RT and choice data of each group. In DDM, we signed momentary sensory evidence accumulated over time to form a decision variable (DV). The accumulation process continues until the DV reaches either the upper or lower bound. The reached bound and the timing of when it reached determined the choice and decision time. Reaction time is modeled as the sum of decision time and additional sensory and motor delays (non-decision time). We fit the DDM to individual behavioral data using maximum-likelihood estimation. Details of this method have been described previously (*35*). The DDM has three free parameters: sensitivity, bound height, and mean non-decision time. The sensitivity *k* determines the linear scaling of the mean momentary evidence in the model with signed stimulus strength. The bound height, *B*, determines the amount of evidence that must be accumulated to reach the upper (+*B*) or lower (− *B*) bound. The nondecision time is drawn from a Gaussian distribution whose mean is a free parameter, and the standard deviation is set to 30% of its mean.

#### Data analysis of Experiment 2

All forces measured during the error clamp trials were transformed into learning coefficients (see the analysis of Experiment 1). In Experiment 2-1, 2-3, and 2-4, the coefficient was calculated based on the force of the strong-force condition to allow direct comparison between the two force conditions. Thus, successful learning in the strong condition results in a coefficient value of 1, and for the weak condition, a coefficient value of 0.5. For Experiment 2-2, since two opposing force fields were used, the coefficients were 1 and -1 for each field.

Learning based on the decision uncertainty context predicts a significant difference in the coefficient between the two fields. However, single-context learning predicted no difference between the two.

#### Comparing the effect across different conditions is Experiment 2

To quantify and compare the effects across the four experiments (Main experiments: 2-1 and 2-1 and control experiments: 2-3 and 2-4), we calculated the expected difference ratio for each experiment using data from the error clamp trials. For example, the maximum expected difference of the learning coefficient in Experiment 2-1, 2-3, and 2-4 will be 0.5 (strong [1] – weak [0.5]). For Experiment 2-2, it was 2 (CW [1] − CCW [-1]). We divided the actual observed difference in the learning coefficient between the two force-field conditions by the maximum expected difference (Fig. 2B).

#### Data analysis of Experiment 3

Data were analyzed in a manner similar to Experiment 2. The analysis was performed separately for the random-dot motion stimuli and arrow sequence stimuli. The correspondence between the contextual effects of the random-dot motion and arrow stimuli was assessed by calculating the expected difference ratio for each stimulus and plotting them against each other (correlation) (Fig. 3E).

### Statistical analysis

For Experiment 1, two-way ANOVA 7 (Group [2] × Coherence level [5]) and the t-test (repeated-measurement) were used for the statistical test. Unless specified otherwise, a t-test (repeated measurement) was used for pair-wise comparison, and the *Bonferroni* method was used to correct multiple comparisons.

## Acknowledgments

The authors are grateful for the help of Ms. Mari Koshimizu in the data collection process and the members of the CiNet Motor Control Unit and HONDA R&D for their helpful insights in our discussions.

## Funding

Part of this work was supported by the Japan Society for the Promotion of Science (Kakenhi:20H00107, 21H00314) and the Japan Science and Technology Agency (ERATO: JPMJER1801).

## Author contributions

Conceptualization: NH and MN

Methodology: KO, AY, GO, MH, and NH

Investigation: KO, AY, GO, MH and NH

Funding acquisition: NH and MN

Project administration: NH and MN

Supervision: NH

Writing – original draft: NH

Writing – review and editing: KO, AY, GO, MN, MH, and NH

## Competing interests

MN is an employee of Honda R&D Co. Ltd. The authors declare that they have no other competing interests.

## Data and materials availability

All data needed to evaluate the conclusion in the paper are present in the paper and Supplementary Materials, and is deposited in OSF website (https://osf.io/n7z4q/). Additional data reported in this paper are available from the corresponding author upon reasonable request.

## Supplementary Materials

Materials and Methods

Supplementary Text (Figure legends)

Figs. S1 to S4

References (33–35)

## Supplementary Materials

### Supplementary Figures & Results

**fig. S1.**
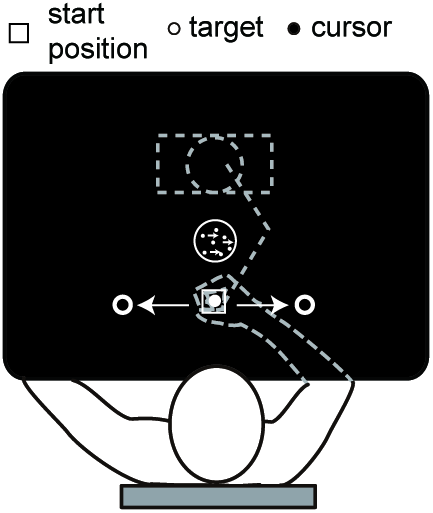
General experimental setup. Participants were seated comfortably in front of a screen placed horizontally in front of them, which prevented the direct vision of their hands. The visual stimulus was presented on the screen using a projector placed above the screen. Participants held a handle of a manipulandum underneath the screen and made a straight reaching movement towards the target (left or right) depending on their perceptual decision.

**fig. S2.**
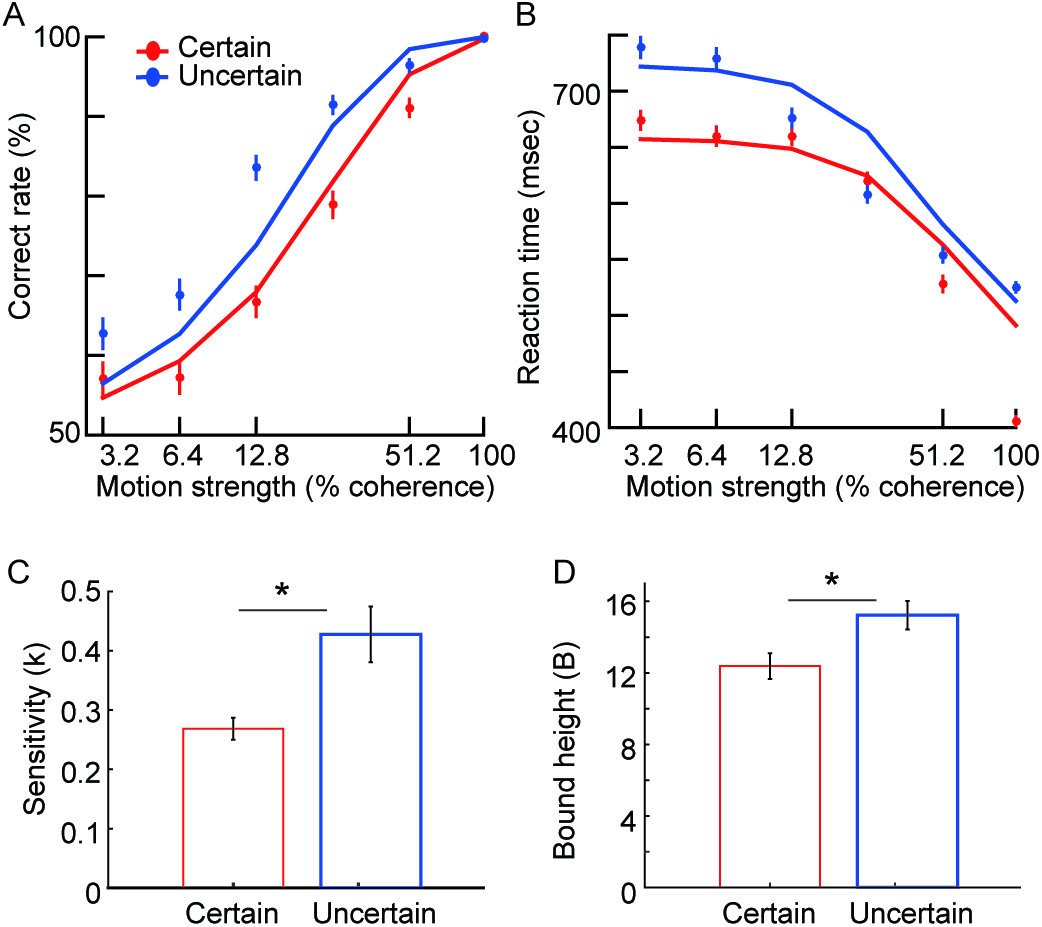
The choice and reaction time data in the retrieval phase of Experiment 1. **A, B**: Correct rate (**A**) and the reaction time (**B**) plotted against different motion coherence levels. A typical psychometric and chronometric function for the random-dot-motion direction decision was observed for both group of participants. Fitted line is derived from the drift-diffusion model parameters applied to the data. **C, D:** Sensitivity to the decision evidence (**C**) and the height of the evidence accumulation bound (**D**) for each Certain and Uncertain group, estimated from the drift-diffusion model (see Supplementary Methods). Uncertain group had significantly higher bound height (**C**) probably because the participants in this group were more cautious in their decision due to the repeated exposure to difficult stimuli. It is likely that difficult stimuli also facilitated perceptual learning in this group and improved their sensitivity *k* (**D**). Error bars indicate the standard error of means across participants. *: *p*<0.05.

**fig. S3.**
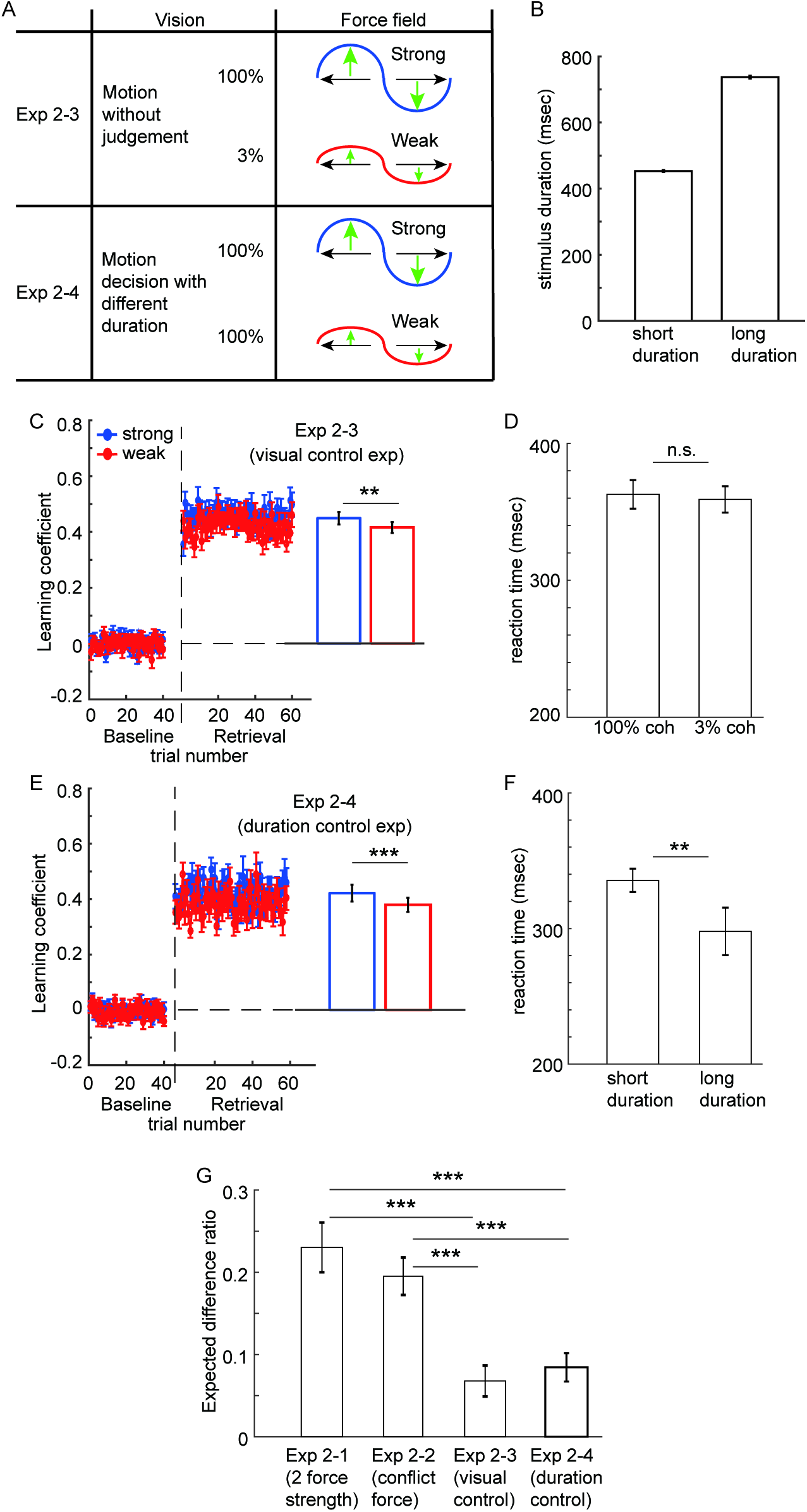
Conditions, stimulus duration, and the results of the control experiments. In Experiment 2-3 (upper panel of **A**), 100% and 3% coherent motion were each associated with strong or weak force fields, but the participants did not make any explicit motion direction decision. Immediately after the disappearance of the motion, the left or right target appeared independent of the motion direction, and the participants reached towards the target. Therefore, the force-field type was only associated with the visual stimulus feature per se, not the uncertainty of the decision. The duration of the visual stimulus was determined by using the reaction time data of Experiment 2-1 (see Supplementary Methods) (**B**). We ensured that the participants paid attention to the motion stimulus by occasionally asking them after the trial which type of stimulus, 100% or 3% coherent motion, was presented (correct rate; 91.4 +- 12.7%). Reaction time, defined as the movement onset from the target presentation (**D**), did not differ depending on the preceding stimulus type, indicating the minimal difference in motor preparation between the two conditions. As shown in **C** and **G**, the effect of learning was only 1/3 of the main experiments (Experiment 2-1, 2-2). In Experiment 2-4 (lower panel of **A**), participants judged the direction of the random-dot motion and learned two different strengths of force fields, but the motion coherence was both fixed at 100%. Here, the two visual conditions differed in the duration of the stimulus (using the same parameter as **B**), but the force-field type was not associated with any difference in the stimulus uncertainty level. The reaction time (**F**) differed between the two conditions, reflecting the difference in the motor preparation level, likely induced by the difference in the foreperiod of action. However, such a difference could not facilitate the learning of the two force fields at the same level as the decision uncertainty context (**E**). The learning effect was again approximately 1/3 that of the main experiments (**G**). Taken together, these control experiments show that decision uncertainty can indeed be a context to tag two different motor memories, which cannot be simply explained by the visual feature or duration of the decision stimulus. Note that **G** is the same Fig. presented in the main text of Fig. 2**E**. Error bars indicate the standard error of the mean across participants. **: *p*<0.01.

**fig. S4.**
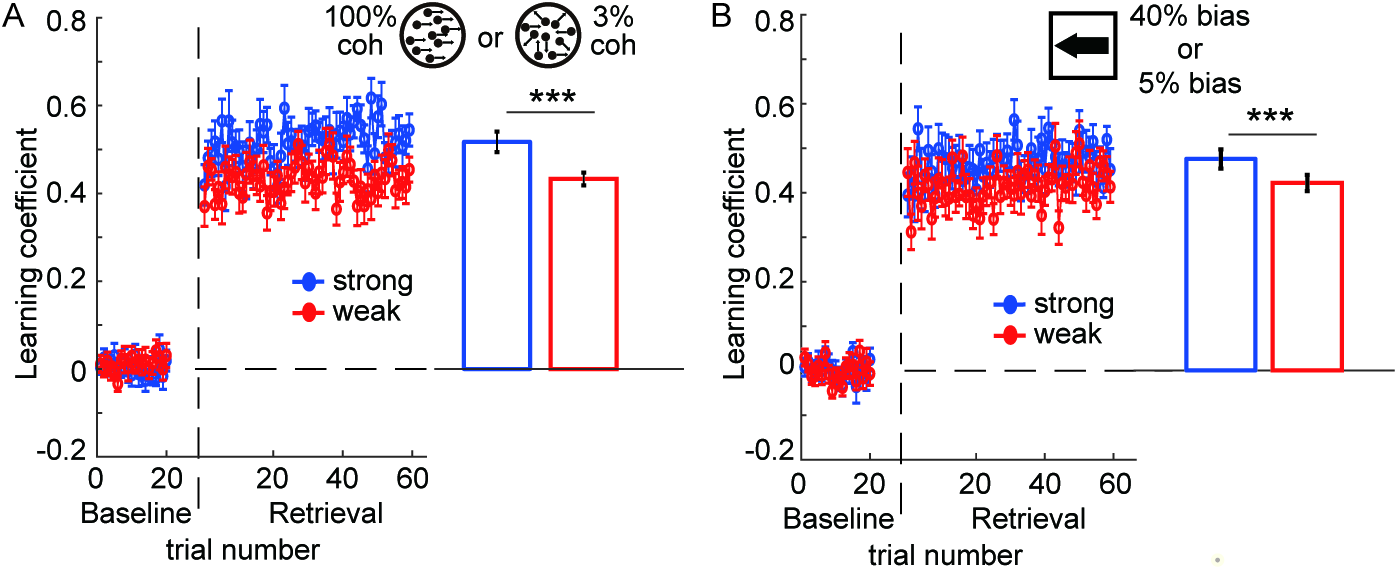
Result of Experiment 3. Radom-dot stimulus; **A**, Arrow sequence; **B**. Note that for the arrow sequence stimuli, participants have never performed the decision in association with any type of force field (see Fig. 3**B** of the main text). Therefore, any difference in force output between different types of arrow stimuli in the retrieval phase is necessary due to the association between the decision uncertainty and the force-field strength leaned through random-dot motion stimuli. Error bars indicate the standard error of means across participants. *: *p*<0.001.

## Notes

### Summary of Updates

Typo is revised

